# Phosphatidylglycerol synthesis facilitates plastid gene expression and light induction of nuclear photosynthetic genes

**DOI:** 10.1101/2021.09.08.459525

**Authors:** Sho Fujii, Koichi Kobayashi, Ying-Chen Lin, Yu-chi Liu, Yuki Nakamura, Hajime Wada

## Abstract

Phosphatidylglycerol (PG) is the only major phospholipid in the thylakoid membrane of chloroplasts. PG is essential for photosynthesis and loss of PG in *Arabidopsis thaliana* results in severe defects of growth and chloroplast development with decreased chlorophyll accumulation, impaired thylakoid formation, and downregulation of photosynthesis-associated genes encoded in nuclear and plastid genomes. However, how the absence of PG affects the gene expression and plant growth remains unclear. To elucidate this mechanism, we investigated the growth and transcriptional profiles of a PG-deficient Arabidopsis mutant *pgp1-2* under various light conditions. Microarray analysis demonstrated that reactive oxygen species-responsive genes were upregulated in *pgp1-2*. Decreased growth light did not alleviated the impaired leaf development and the downregulation of photosynthesis-associated genes in *pgp1-2*, indicating limited impacts of photooxidative stress on the defects of *pgp1-2*. Illumination to dark-adapted *pgp1-2* triggered downregulation of photosynthesis-associated nuclear-encoded genes (PhANGs), while plastid-encoded genes were constantly suppressed. Overexpression of *GOLDEN2-LIKE1* (*GLK1*), a transcription factor regulating chloroplast development, in *pgp1-2* upregulated PhANGs but not plastid-encoded genes along with chlorophyll accumulation. Our data suggest a broad impact of PG biosynthesis on nuclear-encoded genes partially via GLK1 and a specific involvement of this lipid in the plastid gene expression and plant development.

## Introduction

Photosynthesis in chloroplasts provides energy for plant growth but with a simultaneous risk of photooxidative damage. Therefore, the development and functions of chloroplasts are subject to strict regulation in response to various developmental and environmental cues. The expression of photosynthesis-associated nuclear-encoded genes (PhANGs) is strongly affected by the functionality of chloroplasts including tetrapyrrole and carotenoid metabolism, expression of plastid-encoded genes, the redox status of photosynthetic electron transport components, and the homeostasis of reactive oxygen species (Chan *et al.*, 2016). These regulatory mechanisms are known as plastid-to-nucleus retrograde signaling or plastid signaling. GOLDEN2-LIKE (GLK) transcription factors, GLK1 and GLK2 in Arabidopsis (*Arabidopsis thaliana*), are positive regulators of PhANGs, particularly those for chlorophyll biosynthesis and light-harvesting complexes (LHCs), and are involved in the plastid signaling (Kakizaki *et al.*, 2009; Waters *et al.*, 2009; Tokumaru *et al.*, 2017). When the chloroplast functions are severely impaired, the *GLK1* expression is suppressed and the GLK1 protein is degraded, which leads to subsequent downregulation of their target genes including many PhANGs. Consistently, overexpression of GLK1 partially attenuates the downregulation of PhANGs induced by chloroplast dysfunction. Overexpression of GLKs induced ectopic PhANG expression and chlorophyll accumulation in non-photosynthetic organs such as roots, indicating that GLK transcription factors are pivotal regulators of chloroplast biogenesis (Kobayashi *et al.*, 2013*b*).

Expression of plastid-encoded genes is involved in the plastid signaling, as observed in strong downregulation of PhANGs in response to inhibition of plastid gene expression (Chan *et al.*, 2016). Genes in the plastid genome can be classified into three classes; class 1 and class 3 genes are predominantly transcribed by plastid-encoded RNA polymerases (PEP) and nuclear-encoded plastid RNA polymerases (NEP), respectively, whereas class 2 genes are transcribed by both PEP and NEP (Shiina *et al.*, 2005; Pfannschmidt *et al.*, 2015). There are two isoforms of NEPs in Arabidopsis, namely RPOTp localized to plastids and RPOTmp targeted to both plastids and mitochondria. The class 3 genes transcribed by NEP mainly consist of housekeeping genes including *rpo* genes encoding subunits for PEP. Major photosynthesis-associated genes such as *psaA, psbA*, and *rbcL*, which encode core subunits of photosystem I (PSI) and PSII and the large subunit of Rubisco, respectively, are mainly transcribed by PEP and thus classified into the class 1 genes. Similar to bacterial RNA polymerase, PEP requires sigma factors (SIGs), which are encoded in the nuclear genome, for its activity (Shiina *et al.*, 2005; Pfannschmidt *et al.*, 2015). Six *SIG* genes (*SIG1* to *SIG6*) were characterized in Arabidopsis and function in a partially redundant manner. Unlike the other four *SIG*s, disruption of *SIG2* and *SIG6* strongly downregulated the expression of class 1 genes and inhibited chloroplast development in seedlings, indicating a particular importance of these two *SIG*s in chloroplast biogenesis (Woodson *et al.*, 2013). Additional subunits, known as plastid transcriptionally active chromosome proteins (pTACs) or PEP-associated proteins (PAPs), are tightly bound to the PEP complex (Pfalz and Pfannschmidt, 2013; Pfannschmidt *et al.*, 2015). Deficiency of several pTACs or PAPs, such as pTAC2/PAP2, pTAC3/PAP1, or pTAC12/PAP5, strongly blocks the expression of class 1 genes and chloroplast development (Pfalz *et al.*, 2006; Yagi *et al.*, 2012), indicating that at least some pTACs/PAPs are essential for the PEP activity.

Reactive oxygen species (ROS) generated in plastids also function as plastid retrograde signals (Chan *et al.*, 2016). Oxygen molecules that receive electrons from PSI are converted to superoxide and subsequently dismutated to hydrogen peroxide (Asada, 2006). Singlet oxygen is generated by energy transfer from triplet chlorophyll to molecular oxygen or excitation of over-accumulated chlorophyll precursors (Kim and Apel, 2013). Superoxide, hydrogen peroxide, and singlet oxygen affect the expression of the differential gene sets with a partial overlap (op den Camp *et al.*, 2003; Gadjev *et al.*, 2006). Far-red light illumination as well as the loss-of-function mutations of *FLUORESCENCE* (*FLU*) induces the hyperaccumulation of protochlorophyllide in the dark and results in singlet oxygen generation upon light illumination, which causes drastic transcriptomic changes in the nucleus and cell death (Sperling *et al.*, 1997; op den Camp *et al.*, 2003; Page *et al.*, 2016). Deficiency of EXECUTER (EX) 1 and EX2 compromises the upregulation of the stress-induced genes and cell death (Lee *et al.*, 2007). Recently, Page et al. (2016) reported that the mutation of *EX1* and *EX2* partially restores the downregulation of PhANG expression and the impairment of chloroplast differentiation caused by singlet oxygen during deetiolation, indicating the involvement of EXs in the regulation of PhANGs in response to singlet oxygen.

The light reactions of photosynthesis take place in the thylakoid membrane, where glycerolipids provide the lipid bilayer matrix for photosynthetic protein-pigment complexes. The major glycerolipid classes of the thylakoid membrane are glycolipids, and only ∼10 mol% of total thylakoid membrane lipids consist of phospholipids, with phosphatidylglycerol (PG) being major and exclusive phospholipid in plants and cyanobacteria, respectively (Kobayashi, 2016). Despite its relatively low abundance, PG plays crucial roles in photosynthetic electron transport. PG is found in both PSI and PSII (Jordan *et al.*, 2001; Sakurai *et al.*, 2006; Kubota *et al.*, 2010; Umena *et al.*, 2011) and is required for multiple aspects of functions and structures of these photosystem complexes, particularly PSII (Hagio *et al.*, 2000; Sato *et al.*, 2000; Gombos *et al.*, 2002; Sakurai *et al.*, 2003, 2007; Domonkos *et al.*, 2004; Guskov *et al.*, 2009; Kobayashi *et al.*, 2016*a*). Reflecting the essential roles of PG, PG deficiency strongly affects photosynthesis and the growth of both cyanobacteria and plants (Hagio *et al.*, 2000, 2002; Xu *et al.*, 2002; Babiychuk *et al.*, 2003; Kobayashi *et al.*, 2015).

The committed step of PG biosynthesis is the conversion of CDP-diacylglycerol and glycerol-3-phosphate to PG phosphate (PGP) catalyzed by PGP synthase (Kobayashi, 2016). PGP is subsequently dephosphorylated to PG by PGP phosphatase (Lin *et al.*, 2016, 2018; Zhou *et al.*, 2017). Two isoforms of PGP synthase, namely PGP1 and PGP2, have been identified in Arabidopsis (Xu *et al.*, 2002; Hagio *et al.*, 2002; Babiychuk *et al.*, 2003; Tanoue *et al.*, 2014). PGP1 is dually targeted to plastids and mitochondria whereas PGP2 is localized to the ER membrane (Babiychuk *et al.*, 2003; Tanoue *et al.*, 2014). A T-DNA insertion knockout of Arabidopsis PGP1 (*pgp1-2*) caused loss of PG to less than 20% of the wild type, leading to severe impairment of chlorophyll accumulation, thylakoid membrane formation, and photosynthesis (Hagio *et al.*, 2002; Babiychuk *et al.*, 2003; Kobayashi *et al.*, 2015, 2016*a*). In the *pgp1-2* mutant, the expression of PhANGs was remarkably downregulated along with plastid-encoded photosynthetic genes (Kobayashi *et al.*, 2015). However, activation of glycolipid biosynthesis in this mutant by phosphate starvation attenuated the downregulation of these photosynthetic genes with inducing partial development of the thylakoid membrane and chlorophyll accumulation, while the PSII activity was further decreased (Kobayashi *et al.*, 2015). These findings indicate that the expression of photosynthesis genes in both the nucleus and plastids is tightly associated with PG biosynthesis and membrane biogenesis in plastids independently of photosynthetic capability. However, the mechanism of how PG deficiency affects the expression of PhANGs and plastid genes is unclear. To reveal the relationship between plastid PG biosynthesis and expression of photosynthesis genes, we analyzed the transcriptomic profile of *pgp1-2* mutant from various aspects. We also investigated how the expression of the photosynthesis genes responds to the light in absence of plastid PG biosynthesis. Finally, the effects of overexpression of *GLK1* on chloroplast development in *pgp1-2* mutants were analyzed.

## Materials and methods

### Plant materials and growth conditions

The *pgp1-2* (KG10062) (Hagio *et al.*, 2002), *chlh* (SALK_062726) (Huang and Li, 2009) and *ex1* (SALK_002088) (Lee *et al.*, 2007) mutants and the *GLK1*overexpression transgenic line (*GLK1*ox) (Waters *et al.*, 2008) were the Columbia ecotype of Arabidopsis (*Arabidopsis thaliana*). Surface sterilized seeds were vernalized by incubating them in the water at 4°C in darkness. Seeds were germinated and grown on agar-solidified 1× Murashige and Skoog medium containing 3% (w/v) sucrose under continuous white light (∼30 µmol photons m^− 2^ s^− 1^) at 23°C unless otherwise stated. Dark treated seedlings were sampled in the darkroom under the dim green light. Unless otherwise stated, wild type and *chlh* were grown for 14 and 17 d, respectively, and *pgp1-2* and its double lines were grown for 21 d to equalize the growth stages.

### Microarray analysis

Seven-d-old wild-type and 14-d-old *pgp1-2* seedlings were incubated under the continuous white light of 15 µmol photons m^− 2^ s^− 1^ for 7 d. Total RNA was extracted by using the RNeasy Plant Mini Kit (QIAGEN) and its integrity was assessed with the 2100 Bioanalyzer (Agilent Technologies). Arabidopsis Gene Expression Microarray chip version 4 with 43,803 Arabidopsis probes (G2519F; Agilent Technologies) was used for the microarray analysis. Data were analyzed using the GeneSpring GX 14.9 software (Agilent Technologies). Raw data were normalized using the 75th percentile shift for each chip and the signal level of each probe in *pgp1-2* samples was calculated with baseline normalization against the corresponding wild type samples. Features flagged as absent or with an expression value < 100 in all conditions were eliminated from further analyses. Logarithmic fold change values were calculated from three biologically independent replicates and the probes with significant difference (fold change ≥ 2.0 with *P* ≤ 0.05; moderated *t*-test with FDR adjustment according to Benjamini and Hochberg’s method) were selected. To identify the transcript levels, the median of the logarithmic fold change values was calculated from multiple probes annotated to each gene. Data are available from Gene Expression Omnibus (accession no. GSE180205).

### Gene ontology analysis

Gene ontology analysis was performed with online software agriGO v2.0 (http://systemsbiology.cau.edu.cn/agriGOv2/index.php; Tian *et al.*, 2017) using differentially-expressed gene sets as queries. Significant gene ontology terms in the biological process category were represented (*P* < 0.05, Fisher’s exact test with Benjamini-Yekutieli’s correction.)

### Reverse transcription-quantitative PCR analysis

The abundance of mRNA was measured as described (Kobayashi *et al.*, 2015) by using 200 nM gene-specific primers listed in Supplementary Table S1. The relative transcript abundance is presented as the logarithm of the geometric means.

### Chlorophyll determination

Chlorophylls were extracted in 80% (v/v) acetone from shoots crashed in liquid nitrogen. The samples were centrifuged at 10,000 × g for 5 min to remove debris. Chlorophyll concentration was determined as described (Melis *et al.*, 1987) by measuring the absorbance at 720, 663, and 645 nm with Ultrospec 2100 pro spectrophotometer (GE Healthcare).

### Photosynthetic chlorophyll fluorescence analysis

The maximum photosynthetic efficiency of PSII (Fv/Fm) was measured with a JUNIOR-PAM chlorophyll fluorometer (Heinz Walz) as described (Fujii *et al.*, 2014).

### Thiobarbituric acid-reactive substance (TBARS) assay

The 13-d-old wild type and 20-d-old *pgp1-2* seedlings grown under continuous white light (∼30 µmol photons m^− 2^ s^− 1^) were treated with the same white light, strong (∼400 µmol photons m^− 2^ s^− 1^) red light (660 nm LED arrays, ISL-150×150-RHB, CCS, Kyoto), or complete darkness for 1 d. The above-ground parts of seedlings (∼0.2 g fresh weight) were ground in liquid nitrogen and homogenized in a 1-mL extraction buffer (175 mM NaCl in 50 mM Tris-HCl, pH 8.0). The homogenate was equally divided into two aliquots, one of which was mixed with 0.5 mL of 0.5% (w/v) thiobarbituric acid (TBA) in 20% (w/v) trichloroacetic acid (TCA) for the TBA reaction, and the other was mixed with 0.5 mL of 20% (w/v) TCA to prepare the background control. Both samples were heated at 95°C for 25 min, followed by centrifugation at 10,000 × g for 20 min to remove cell debris. The absorbance of the supernatant was measured at 532 nm, with the absorbance at 600 nm subtracted to account for non-specific turbidity. After subtracting the absorbance of the background control, the amount of malonaldehyde was calculated using an extinction coefficient of 155 mM^− 1^ cm^− 1^ (Larkindale and Knight, 2002).

### Trypan blue staining

Whole seedlings of 14-d-old wild type and 21-d-old *pgp1-2* were immersed in 10 mL of ethanol-lactophenol (2 volumes of ethanol and 1 volume of phenol-glycerol-lactic acid-water (1:1:1:1, w/w/w/w)) containing 0.05% (w/v) trypan blue. The seedlings were then boiled for 10 min and kept at room temperature for 30 min. After washing the seedlings with clearing solution (2.5 g chloral hydrate in 1 mL deionized water) overnight twice, the samples were suspended in 50% (v/v) glycerol and observed under a stereomicroscope (MZ 16FA; Leica) and a color CCD camera (VB-7010; Keyence). The relative intensity of trypan blue staining was analyzed with Image J software.

## Results

### Reactive oxygen species-inducible genes are upregulated in *pgp1-2*

To understand how the loss of plastid PG biosynthesis affects seedling growth, we compared the transcriptome profile between *pgp1-2* and wild-type seedlings by microarray analysis. Out of 43,603 probes annotated to the Arabidopsis genes in the microarray, we identified 12,400 probes with statistically different signal intensity between *pgp1-2* and wild type (*P*-value ≤ 0.05). Among these probes, 2008 probes were upregulated ≥ 2-fold and 2555 probes were downregulated ≤ 0.5-fold in *pgp1-2* compared with wild type. Almost all of the probes annotated to the plastid genome were found in the downregulated group in *pgp1-2*, whereas those for mitochondrial genome were mainly found in the upregulated group (Figure 1). Gene ontology analysis of nuclear-encoded genes showed that stress response genes were enriched in the upregulated gene group, whereas genes involved in photosynthesis were enriched in the downregulated gene group (Table 1).

**Figure 1.**
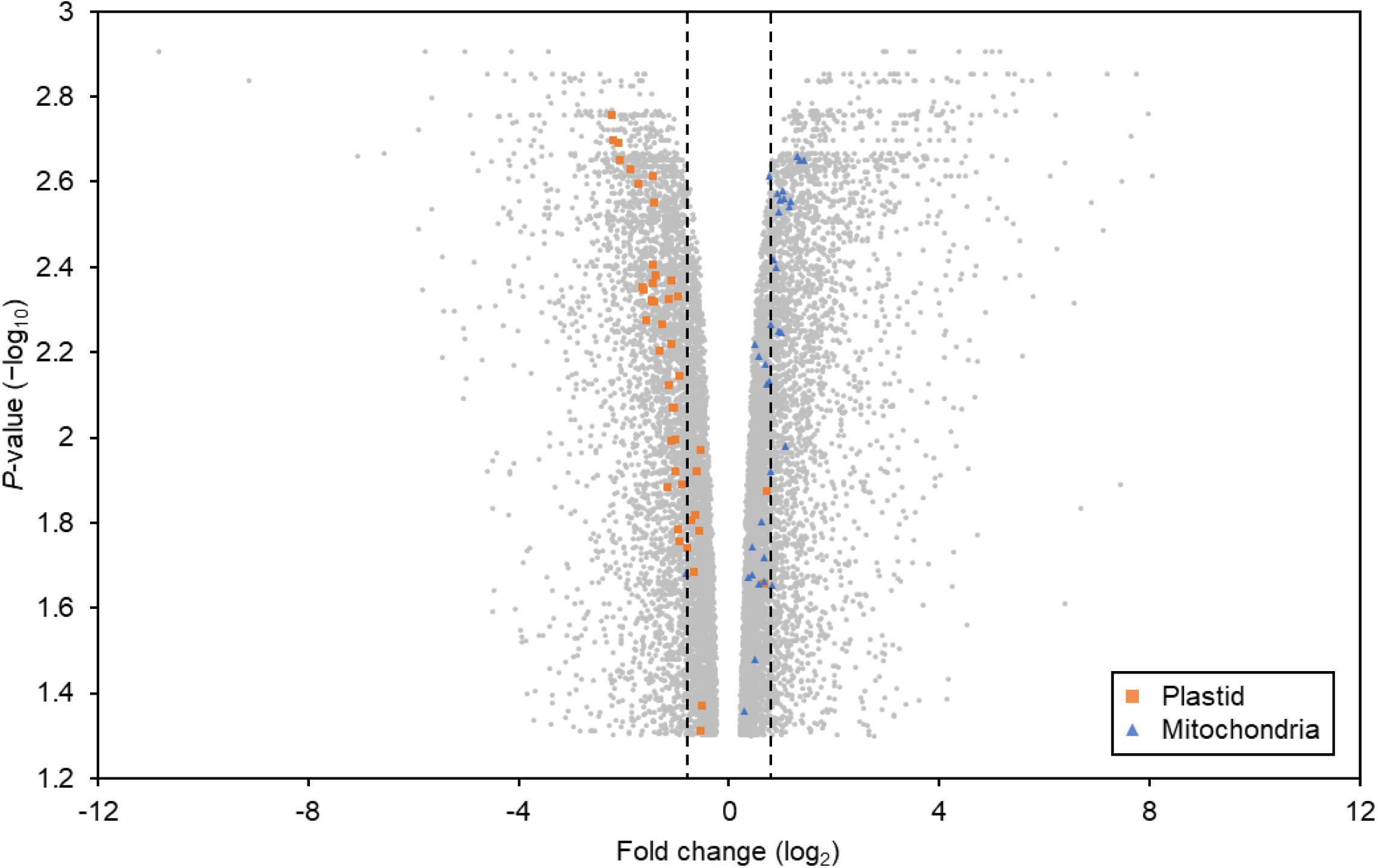
Volcano plot of the microarray probes represented statistically differential signal intensities between *pgp1-2* and wild type (*P* ≤ 0.05). Organellar-encoded genes for proteins and ribosomal RNA are represented. Vertical dashed lines indicate 2-fold and 0.5-fold changes.

**Table 1.** Enrichment of GO terms in differentially expressed genes in *pgp1-2*. Top 10 categories are represented.

| | GO term | Description | *P*-value ($-\log_{10}$) |
| --- | --- | --- | --- |
| **Downregulated** | GO:0015979 | photosynthesis | 63.47 |
|  | GO:0019684 | photosynthesis, light reaction | 38.14 |
|  | GO:0006091 | generation of precursor metabolites and energy | 30.57 |
|  | GO:0050896 | response to stimulus | 25.70 |
|  | GO:0055114 | oxidation-reduction process | 24.64 |
|  | GO:0042221 | response to chemical | 22.48 |
|  | GO:0044710 | single-organism metabolic process | 21.38 |
|  | GO:0009767 | photosynthetic electron transport chain | 20.11 |
|  | GO:0010033 | response to organic substance | 19.70 |
| **Upregulated** | GO:0050896 | response to stimulus | 9.59 |
|  | GO:0042221 | response to chemical | 7.80 |
|  | GO:0006950 | response to stress | 7.23 |
|  | GO:0006855 | drug transmembrane transport | 6.36 |
|  | GO:0042493 | response to drug | 6.00 |
|  | GO:0015893 | drug transport | 6.06 |
|  | GO:0009607 | response to biotic stimulus | 5.12 |
|  | GO:0051704 | multi-organism process | 5.10 |
|  | GO:0097659 | nucleic acid-templated transcription | 5.03 |

From this result, we hypothesized that dysfunction of photosynthetic machinery caused by PG deficiency leads to production of ROS and induces the expression of stress-responsive genes. To test this hypothesis, we investigated the mRNA levels of genes specifically induced by singlet oxygen, superoxide, or hydrogen peroxide (Gadjev *et al.*, 2006). In *pgp1-2*, 9.1%, 11%, and 14% of genes specific to singlet oxygen, superoxide, and hydrogen peroxide, respectively, were upregulated more than twofold compared with the wild-type levels (Fig. 2A, Supplementary Table S2). 15%, 23% and 29% of each subset of ROS-responsive genes were upregulated statistically significantly (*P* ≤ 0.05) in the mutant. This result indicates that hydrogen peroxide-inducible genes were the most preferentially upregulated in *pgp1-2* among three subsets of ROS-inducible genes.

**Fig. 2.**
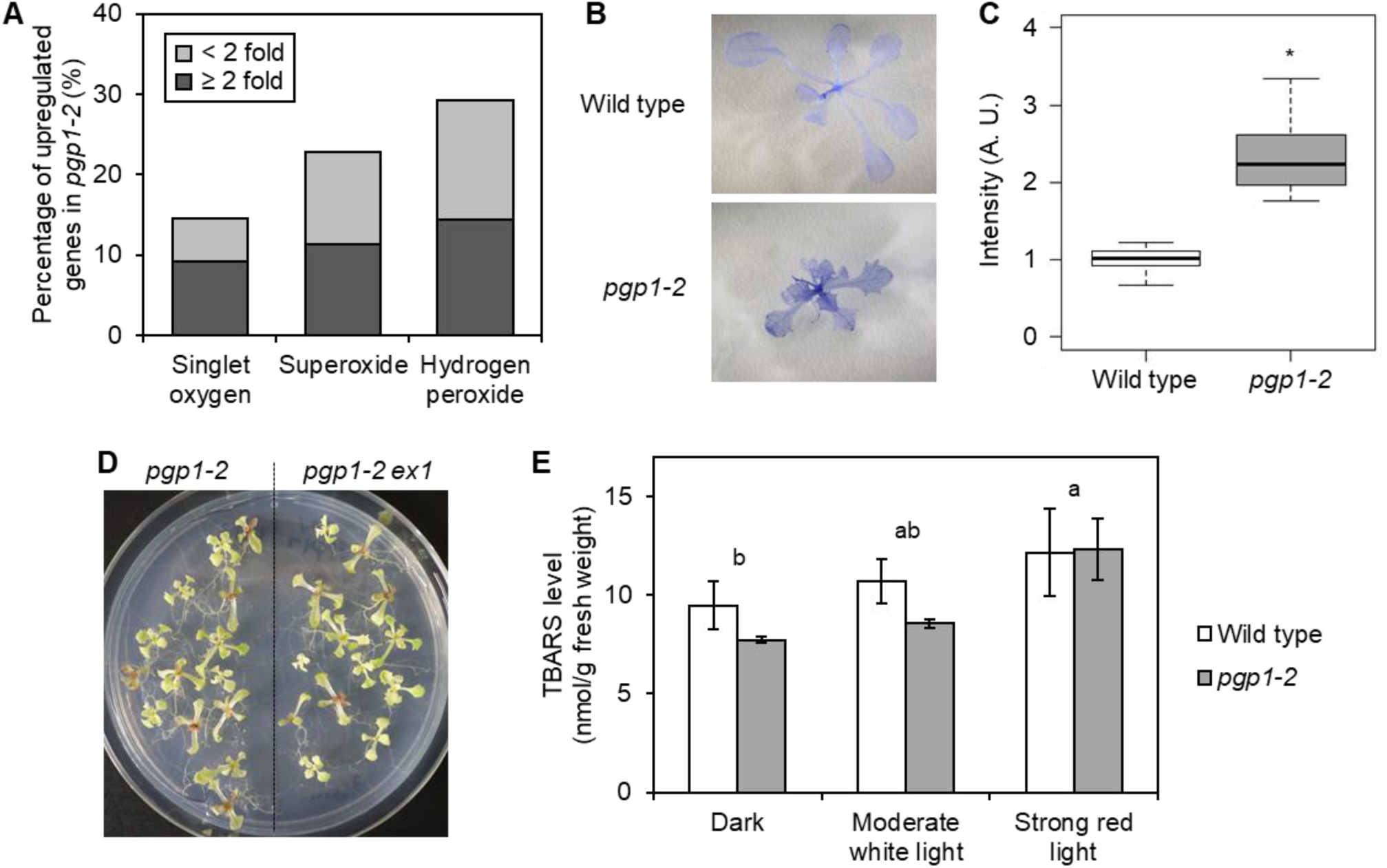
Oxidative stress in *pgp1-2*. (A) Percentages of the number of upregulated genes in *pgp1-2* among ROS-inducible genes. (B) Trypan blue staining of wild-type and *pgp1-2* seedlings. Representative data from more than 10 biological replicates were shown. (C) Quantitative analysis of trypan blue staining. The relative intensity of the trypan blue signal was shown (the average in the wild type = 1). Data are the distribution of 12 (wild type) and 13 (*pgp1-2*) biological replicates. An asterisk indicates the statistical significance (*P* < 0.05, Welch’s *t*-test). (D) Phenotype of 21-d-old *pgp1-2 ex1* double knockout mutant. (E) Thiobarbituric acid reactive substance (TBARS)-assay in wild-type and *pgp1-2* seedlings. Seedlings were incubated in darkness or under moderate white light of 30 µmol photons m^− 2^ s^− 1^ or strong red light of 400 µmol photons m^− 2^ s^− 1^ for 1 d before sampling. Levels of TBARS are indicated as means ± SE from three biological replicates. Different letters indicate significant differences (*P* < 0.05, Tukey-Kramer multiple comparison test after two-way ANOVA). There were no significant effect of genotypes and no significant interaction between light conditions and genotypes.

To assess whether the *pgp1-2* mutation causes cell death in leaves during seedling development, we stained *pgp1-2* and wild-type seedlings with trypan blue, which visualizes dead cells. The mutant leaves showed dark blue staining in contrast to faint staining in wild type, indicating enhanced cell death in *pgp1-2* (Fig. 2B, C). EXECUTER1 (EX1) and EX2 play central roles in singlet oxygen-mediated cell death and downregulation of photosynthesis-related genes (Lee *et al.*, 2007; Page *et al.*, 2016). To assess the contribution of EXs to the severe phenotype of *pgp1-2*, we crossed the *pgp1-2* with *ex1* knockout lines and obtained the double knockout mutant *pgp1-2 ex1*. The phenotype of *pgp1-2 ex1* was not distinguishable from that of *pgp1-2*, indicating that EX1-mediated signal transduction is not a major pathway to cause the phenotype of *pgp1-2* (Fig. 2D).

To test whether oxidative stress was increased in *pgp1-2*, we measured the level of TBARS, which is known to accumulate in response to lipid peroxidation by ROS (Larkindale and Knight, 2002). Because strong red light caused photoinhibition in *pgp1-2*, but not in wild type, presumably due to the dysfunction of PSII at the donor side in the mutant (Kobayashi *et al.*, 2016*a*), we compared TBARS levels in seedlings treated with relatively strong red light (400 µmol photons m^− 2^ s^− 1^) for 24 h with those treated with moderate white light (∼30 µmol photons m^− 2^ s^− 1^) and darkness. However, no significant difference was observed in TBARS levels between wild type and the mutant on a fresh weight basis under all light conditions examined (Fig. 2E).

### Expression of photosynthesis-related genes is drastically changed in *pgp1-2*

Our transcriptome analysis revealed that more than 70% of the genes involved in photosynthetic electron transport were downregulated in *pgp1-2* compared to the wild type, whereas none of them were upregulated. Most of the genes encoding subunits for photosystems, antenna complexes, the cytochrome *b*_6_*f* complex, and the ATP synthase showed significantly decreased mRNA levels in *pgp1-2* (Fig. 3, Supplementary Table S3). Downregulation was particularly strong in core subunits of LHCII, namely *LHCB1* and *LHCB2*. In addition, genes encoding plastocyanin and those for major ferredoxin and ferredoxin-NADP^+^ reductase (FNR) in photosynthetic tissues were suppressed in *pgp1-2*. Although half of the core genes of NAD(P)H:plastoquinone dehydrogenase (NDH) complex, which are plastid-encoded, were not largely affected by PG deficiency, all of the peripheral NDH subunit genes encoded in the nucleus were downregulated more than twofold. Genes for the major isoforms of the Calvin-Benson cycle enzymes (Stitt *et al.*, 2010) were also downregulated in *pgp1-2* (Fig. 3). These results show that PG deficiency has a broadly suppressive effect on the transcript level of both stromal and membranous photosynthetic components in chloroplasts.

**Fig. 3.**
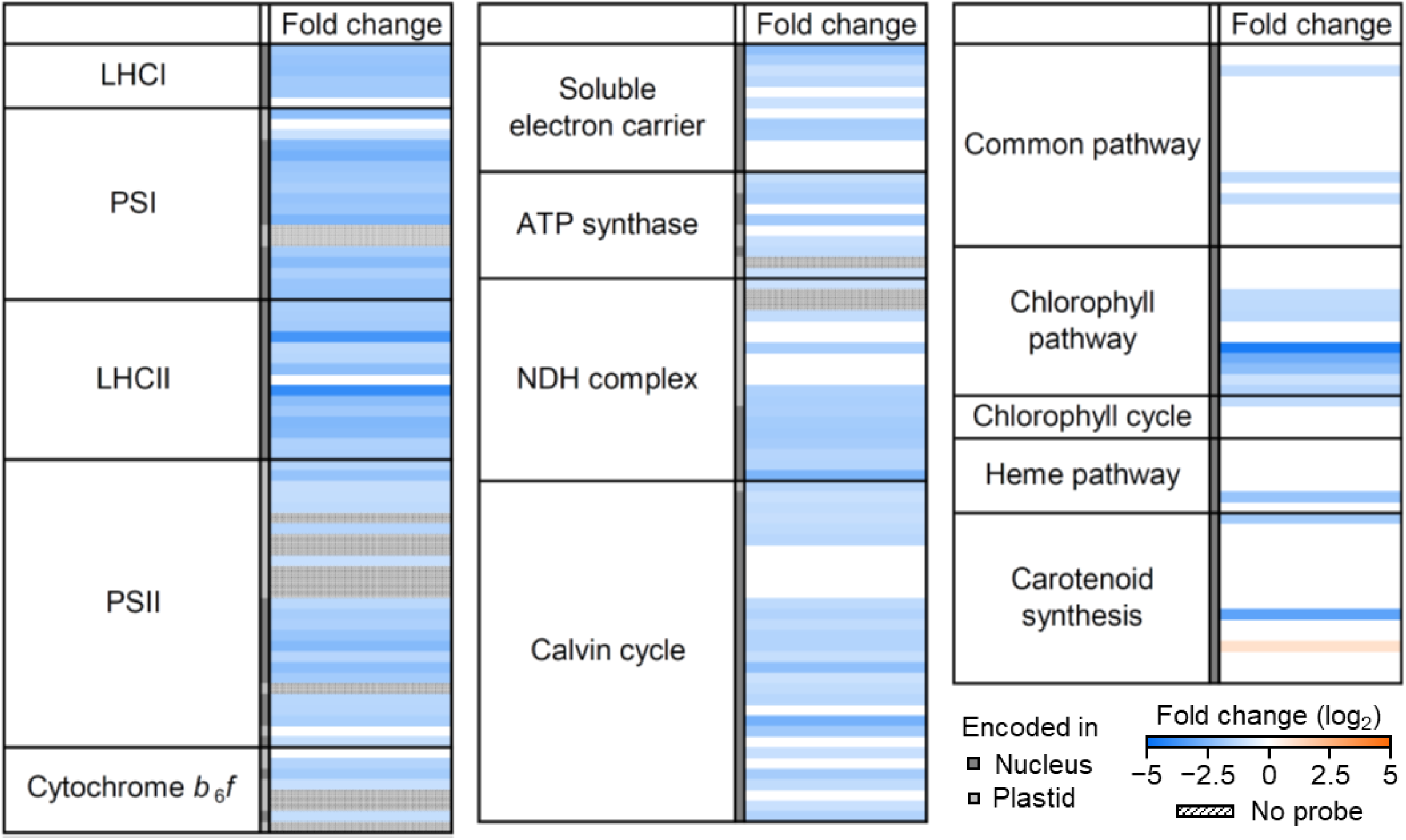
Microarray analysis of photosynthesis-associated genes in *pgp1-2*. Logarithmic fold change between *pgp1-2* and the wild type is shown. Data are means from three biological replicates. Median values are represented for genes with multiple probes. Changes larger than 2-fold or smaller than 0.5-fold are indicated with colors. For more details, see Supplementary Table S3.

In the tetrapyrrole metabolism pathway, some genes specifically involved in chlorophyll biosynthesis were significantly downregulated in *pgp1-2*, whereas genes for the heme biosynthesis pathway were not largely affected (Fig. 3). Particularly, the transcript levels of all *POR* genes (*PORA, PORB*, and *PORC*) encoding isoforms of light-dependent NADPH:protochlorophyllide oxidoreductase were strongly decreased in the mutant. On the other hand, PG deficiency had limited effects on the transcript accumulation of genes involved in carotenoid biosynthesis.

### PEP-dependent expression of plastid-encoded genes was suppressed in *pgp1-2*

To understand how the photosynthetic gene expression in plastids was downregulated in the *pgp1-2* mutant, we characterized the transcript profiles of protein- and rRNA-coding genes in the plastid genome in more detail (Fig. 4A). All of 20 probed class 1 genes were downregulated in the mutant and 13 of them were decreased to more than 0.5-fold of the wild-type level. Some of the class 2 genes, including genes for ATP synthase subunits, but no class 3 genes, were also downregulated below 0.5 of the wild-type level in this mutant. This result indicates that PEP-dependent gene expression is preferentially suppressed in *pgp1-2*. Therefore, we next examined the transcript level of genes involved in PEP-dependent plastid gene expression (Fig. 4B). The expression of genes encoding PEP core components, *rpoA, rpoB, rpoC1*, and *rpoC2*, was not significantly changed in *pgp1-2* compared with wild type. Among six sigma factors that facilitate the PEP activity in Arabidopsis, *SIG1* and *SIG4* were downregulated in *pgp1-2* to 30% of the wild-type level. Besides sigma factors, numbers of nucleoid-associated proteins such as pTAC and PAP proteins are required for PEP activity (Pfalz and Pfannschmidt, 2013; Pfannschmidt *et al.*, 2015). The mRNA levels of nucleoid-associated genes, namely *pTAC16, pTAC8*/*CURT1B, MFP1*, and *RAP* were significantly decreased in the *pgp1-2* mutant (Fig. 4C). In addition, mRNA levels of genes encoding sedoheptulose-1,7-bisphosphatase (SBPase), ATP synthase subunits, the large subunit of Rubisco, Rubisco activase (RCA), calcium-sensing receptor (CAS), and serine/threonine-protein kinase 8 (STN8) which are found in plastid nucleoid fraction (Melonek *et al.*, 2016), were also downregulated in *pgp1-2*. As for NEPs, the expression level of *RPOTp* was not significantly changed and transcripts of *RPOTmp* were even three times higher in *pgp1-2* than the wild type (Fig. 4B).

**Fig. 4.**
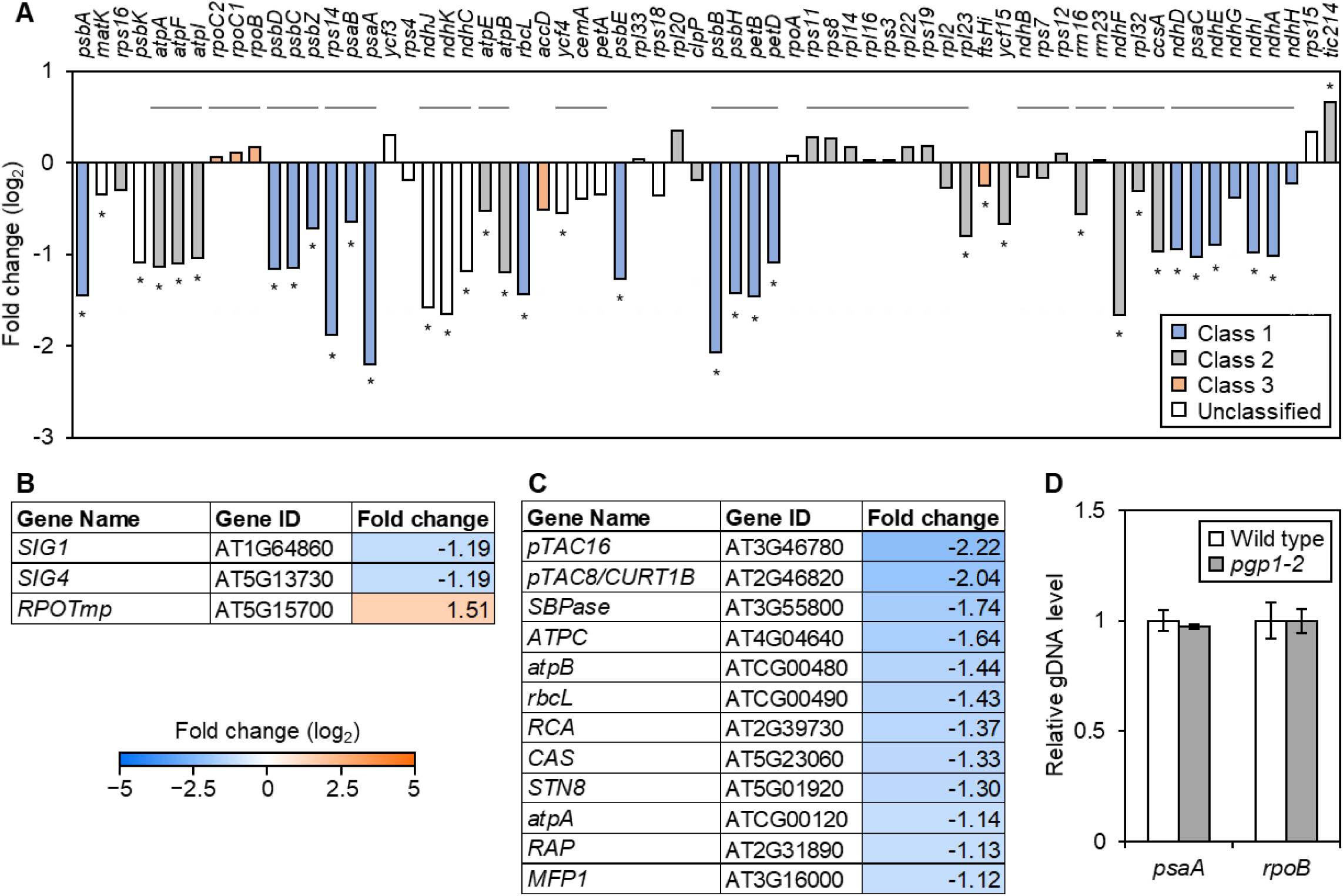
Expression of plastid-encoded genes in *pgp1-2*. (A) Changes in the transcript level of plastid-encoded genes probed in the microarray. Gray lines indicate the operons. Asterisks indicate statistical significance (*P* < 0.05). Data are geometric means from three biological replicates. (B) and (C), List of differentially expressed genes encoding core components of plastid RNA polymerase (B) and nucleoid-associated protein in plastids (C). Data are means from three biological replicates. In (A) to (C), logarithmic fold change between *pgp1-2* and the wild type is shown. (D) Relative content of plastid DNA in the wild type and *pgp1-2*. DNA levels of two loci of the plastid genome were quantified by real-time PCR, normalized to the nuclear DNA level of the *ACTIN8* locus, and presented as the difference from the wild-type level. Data are means ± SE from three biological replicates. No significant differences were detected between the two genotypes (*P* > 0.05, Student’s *t*-test).

Since the copy number of the plastid genome may also affect the transcript level in plastids, we assessed the relative amount of plastid DNA by quantitative PCR. In the two regions we tested, the copy number of the plastid genome was similar between *pgp1-2* and the wild type (Fig. 4D).

### mRNA accumulation of photosynthesis-associated genes was suppressed in *pgp1-2* regardless of light intensity

To test whether the severe impairment of leaf development and downregulation of photosynthesis-associated genes in the *pgp1-2* mutant were caused as a response to the light stress, we grew the mutant and wild-type seedlings under different light intensities ranging from 5 to 75 µmol photons m^− 2^ s^− 1^. The color phenotype of *pgp1-2* mutants was not largely affected by the different light intensities and the impairment of leaf development was not ameliorated under the low light condition (Fig. 5A).

**Fig. 5.**
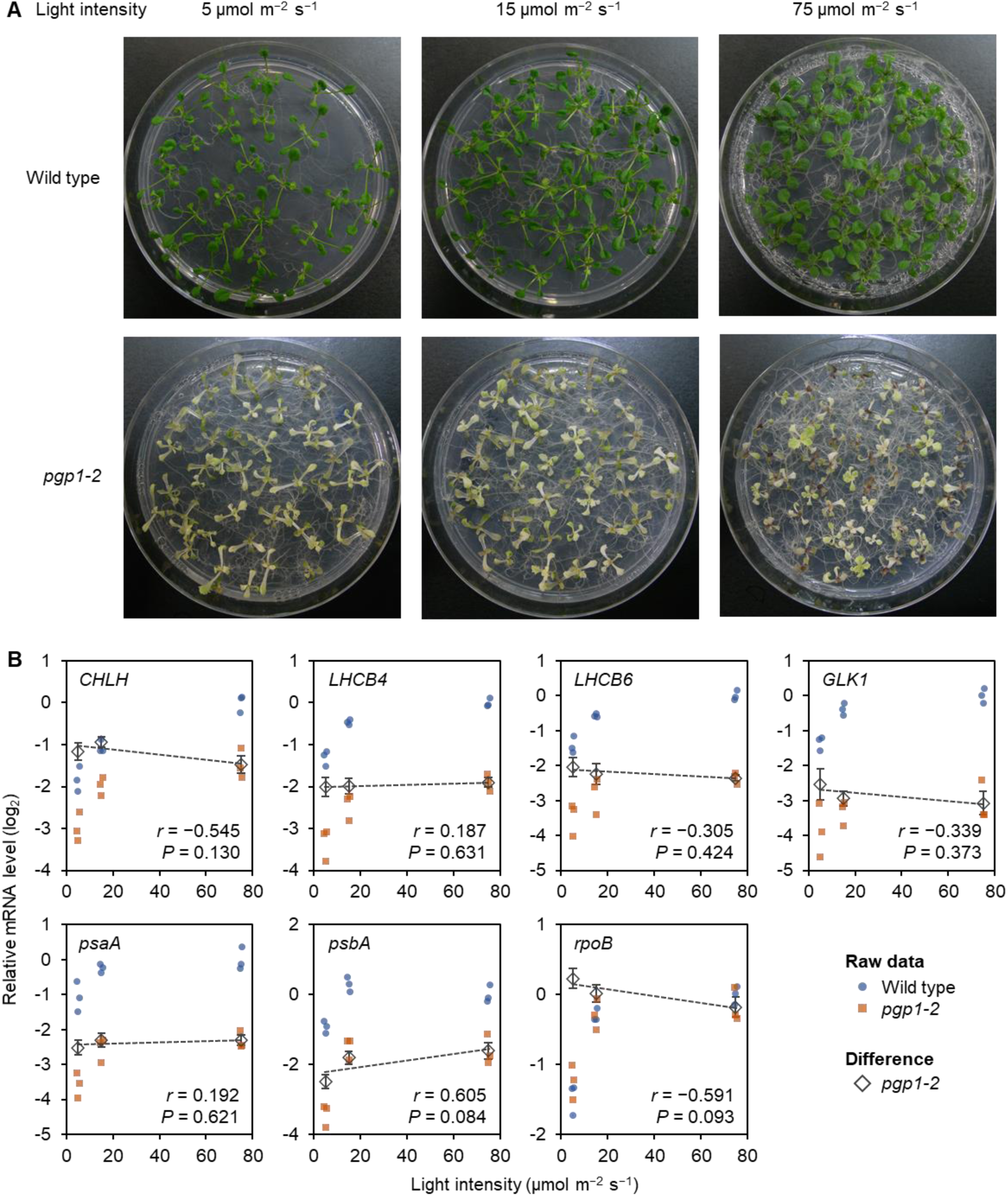
Expression of photosynthesis-related genes in *pgp1-2* grown under different light conditions. (A) 7-d-old wild type and 14-d-old *pgp1-2* seedlings were grown under indicated light intensities for 7 d. (B) Transcript levels of photosynthesis-associated genes in seedlings under different light intensities. Transcript levels are normalized to *ACTIN8* and presented as the difference from the wild type under 75 µmol photons m^− 2^ s^− 1^. Gray diamonds indicate the transcript levels in *pgp1-2* normalized to the mean of wild-type transcript levels in each light condition. Data are means ± SE from three biological replicates. Correlation coefficients (*r*) of the normalized transcript levels and their *P*-values are indicated.

To understand the light response of PhANGs in the mutant, we carried out reverse transcription-quantitative PCR (RT-qPCR) analysis of mRNA levels of *LHCB4* and *LHCB6* encoding LHCII subunits, *CHLH* encoding the H subunit of Mg-chelatase for chlorophyll biosynthesis, and *GLK1*. To characterize plastid gene expression, we determined mRNA levels of *rpoB* encoding the β subunit of PEP, which is transcribed by NEP, in addition to PEP-dependent genes *psaA* and *psbA*. Under 75 µmol photons m^− 2^ s^− 1^ light, mRNA levels of all tested genes except for *rpoB* were higher in wild type than *pgp1-2* (Fig. 5B). Decreased light intensity to 15 and further to 5 µmol photons m^− 2^ s^− 1^ strongly suppressed mRNA accumulation of all tested genes both in wild type and the mutant. As a result, the mutant-to-wild type ratios of the mRNA levels of these genes did not show significant correlation with light intensity, indicating that the ratios were almost constant across the different light intensities. Of note, the RT-qPCR analysis revealed constantly lower mRNA levels of *CHLH* in *pgp1-2* than in wild type, although the microarray analysis did not show significant changes of this gene in this mutant, which would reflect the higher detection capability of RT-qPCR analysis for a targeted gene than the large-scale microarray analysis.

### Dark incubation partially attenuated downregulation of photosynthesis genes in *pgp1-2*

To further evaluate the contribution of light to the suppression of photosynthetic genes in *pgp1-2*, we investigated the mRNA levels of PhANGs and plastid-encoded genes after transferring the seedlings to complete darkness (Fig. 6). In addition to *LHCB6, CHLH*, and *GLK1*, we tested the transcript levels of four PhANGs, namely *LHCA4* encoding one of LHCI subunits, *HEMA1* encoding the major isoform of glutamyl-tRNA reductase for chlorophyll and heme biosynthesis, and *SIG1* and *SIG4*. As for class 1 plastid-encoded genes, the mRNA levels of *psaA, psbA, rbcL, petB* encoding the *b*_6_ subunit of the cytochrome *b*_6_*f* complex, *ndhA* encoding one of the core subunits of the NDH complex, and *rps14* encoding the S14 subunit of plastid ribosome were analyzed. Transcript accumulation of two class 3 genes, namely *accD* encoding the β-carboxyl transferase subunit of acetyl-CoA carboxylase and *rpoB* was also tested. In wild type, dark incubation for 6 h greatly decreased the mRNA levels of all genes except *rps14*. Prolonged dark treatment for 1 and 7 d further decreased the mRNA levels of these genes except for *accD*. Similarly, in *pgp1-2*, the dark treatment decreased mRNA levels of photosynthetic genes except for *GLK1*, whose mRNA level was doubled after 24 h-dark incubation. However, in *pgp1-2*, the dark-induced downregulation of these genes was less pronounced than the wild type, so the *pgp1-2*-to-wild type ratio of mRNA levels increased in many genes by the dark treatment for 24 h. However, prolonged dark treatment for 7d did not further increase the *pgp1-2*-to-wild type ratio in most of the genes tested.

**Fig. 6.**
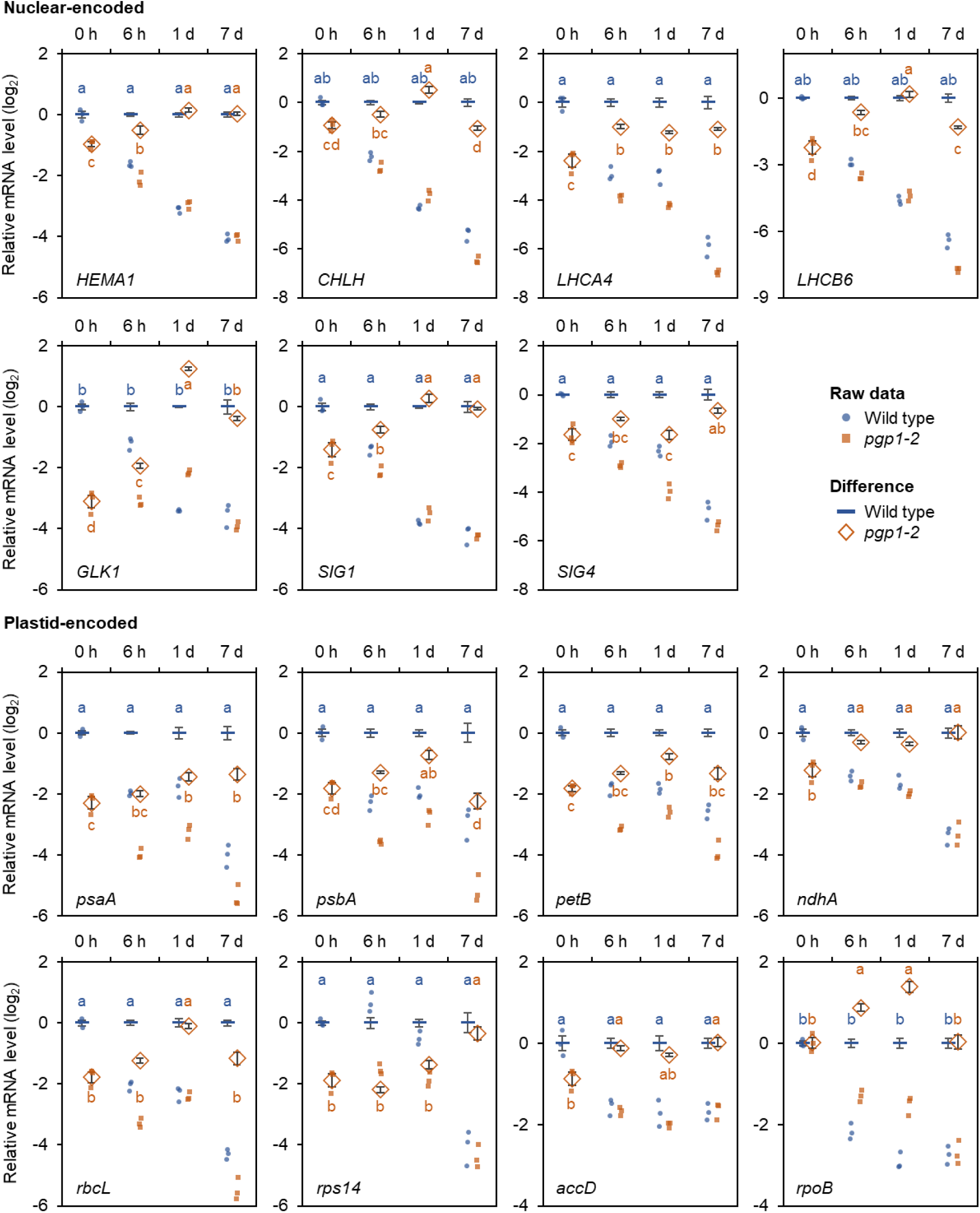
Effect of dark treatment on expression of photosynthesis-related genes in *pgp1-2*. Transcript levels of photosynthesis-associated genes encoded in the nucleus and plastids are shown. Seedlings of 14-d-old wild type and 21-d-old *pgp1-2*, which were incubated under continuous light (0 h) or incubated in the dark for 6 h, 1 d, and 7 d before sampling, were used for the experiments. Transcript levels are normalized to *ACTIN8* and presented as the difference from the wild type without dark treatment (closed circles and squares). Then the mRNA level of each gene was normalized to the mean value of the corresponding gene in wild type (horizontal bars and open diamonds). Data are means ± SE from three biological replicates. Different letters indicate significant differences (*P* < 0.05, Tukey-Kramer multiple comparison test after two-way ANOVA).

### Light-dependent induction of photosynthesis gene expression was severely inhibited in *pgp1-2*

To reveal how the *pgp1-2* mutation affects the light-dependent induction of photosynthesis genes, we re-illuminated the plants after 24 h dark treatment and analyzed temporal changes in mRNA levels of nuclear-encoded *HEMA1, CHLH*, and *LHCB6* and plastid-encoded *psbA* and *rbcL* (Fig. 7). In wild type, the mRNA levels of the nuclear-encoded genes rapidly increased to more than 10-fold with 3 h of illumination, whereas those of plastid-encoded genes gradually increased from 1 h to 12 h of illumination and became 3 times higher than those in dark-incubated plants after 12-h illumination. In the *pgp1-2* mutant, mRNA levels of the nuclear-encoded genes increased in response to light until 3 h of illumination, but to a much lower extent than those in wild type. The transcript levels of *CHLH* and *LHCB6* gradually decreased from 3 h to 12 h of illumination in the mutant. The transcript levels plastid-encoded genes were also induced from 1 h to 6 h of illumination in *pgp1-2* but did not further increased after 6 h. Moreover, the mutant showed the pronounced decrease of mRNA levels in the first hour of illumination, particularly in *rbcL*. Consequently, the maximal levels of *psbA* and *rbcL* after illumination were less than two and one fold of their levels before illumination, respectively.

**Fig. 7.**
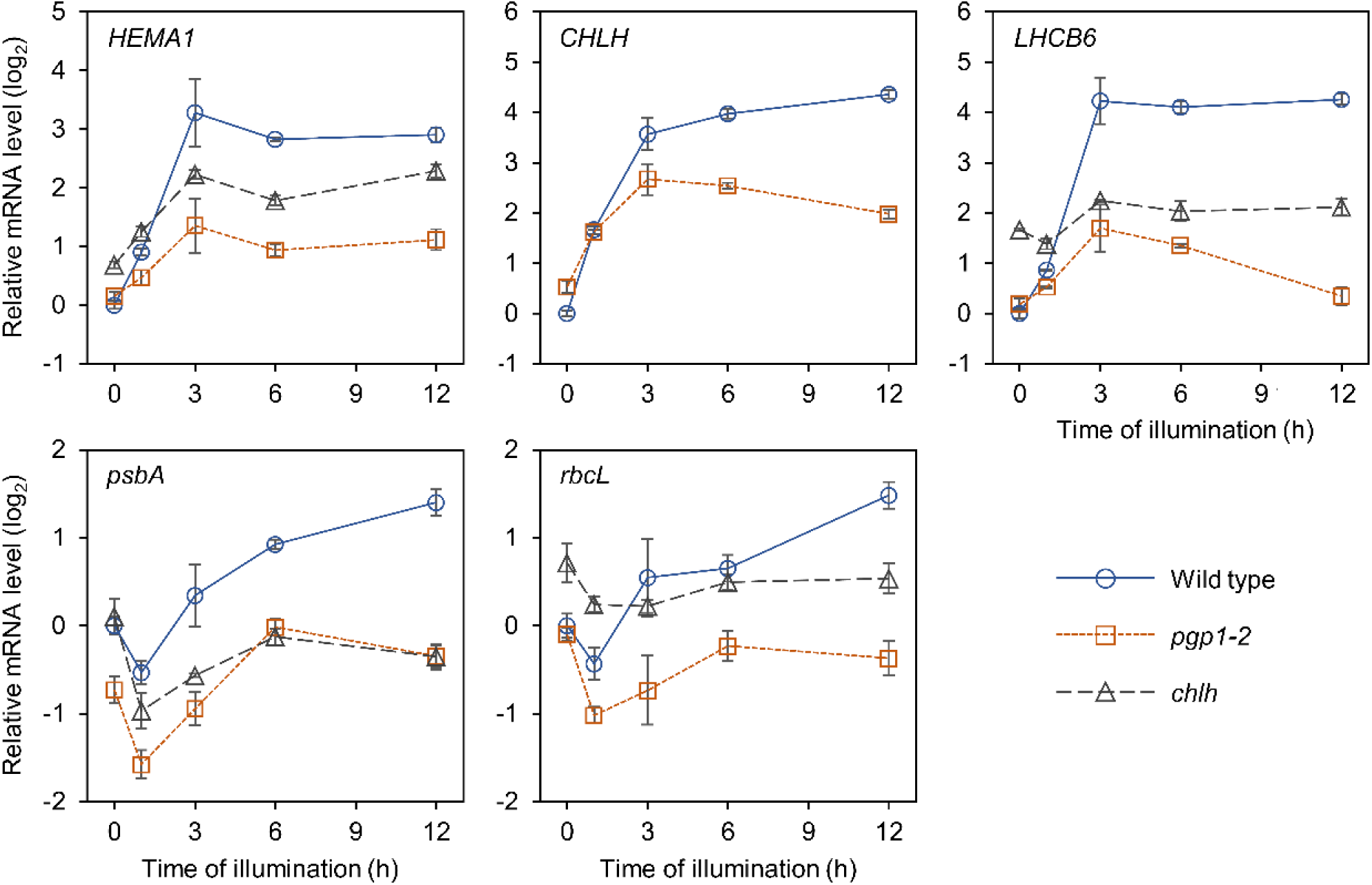
Light-dependent induction of photosynthesis-associated genes in *pgp1-2* and *chlh*. Seedlings were grown for 13 d (wild type), 20 d (*pgp1-2*), and 16 d (*chlh*) under continuous light, followed by the dark treatment for 24 h. Then seedlings were illuminated for 12 h under continuous light. Transcript levels were normalized to *ACTIN8* and presented as the difference from the wild type before illumination. Data are means ± SE from three biological replicates.

To test if the suppression pattern of photosynthesis-associated genes observed in this study is specific to the PG deficiency, we analyzed the transcriptional profile of these genes in the chlorophyll deficient mutant *chlh*, which carries a T-DNA insertion in the coding region of *CHLH* (Huang and Li, 2009). Although the *chlh* mutant showed retarded growth and yellowish leaf color due to the complete loss of chlorophylls, the morphology of the leaves was not as disordered as those of *pgp1-2* (Fig. 8A). As in the *pgp1-2* mutant, the steady-state mRNA levels of both nuclear- and plastid-encoded genes under continuous light were much lower in *chlh* than in the wild type (Fig. 8B). Although the mRNA levels of *HEMA1* and *LHCB6* in *chlh* decreased after dark treatment for 24 h, their levels were significantly higher than those in dark-treated wild type. In *chlh*, the increase of the *GLK1* transcript level by dark treatment was more pronounced than that in *pgp1-2*. Unlike in wild type, the mRNA level of plastid-encoded genes was not largely changed by dark treatment in *chlh*, which led to a significant increase of the *chlh*-to-wild type ratio in all investigated genes (Fig. 8B).

**Fig. 8.**
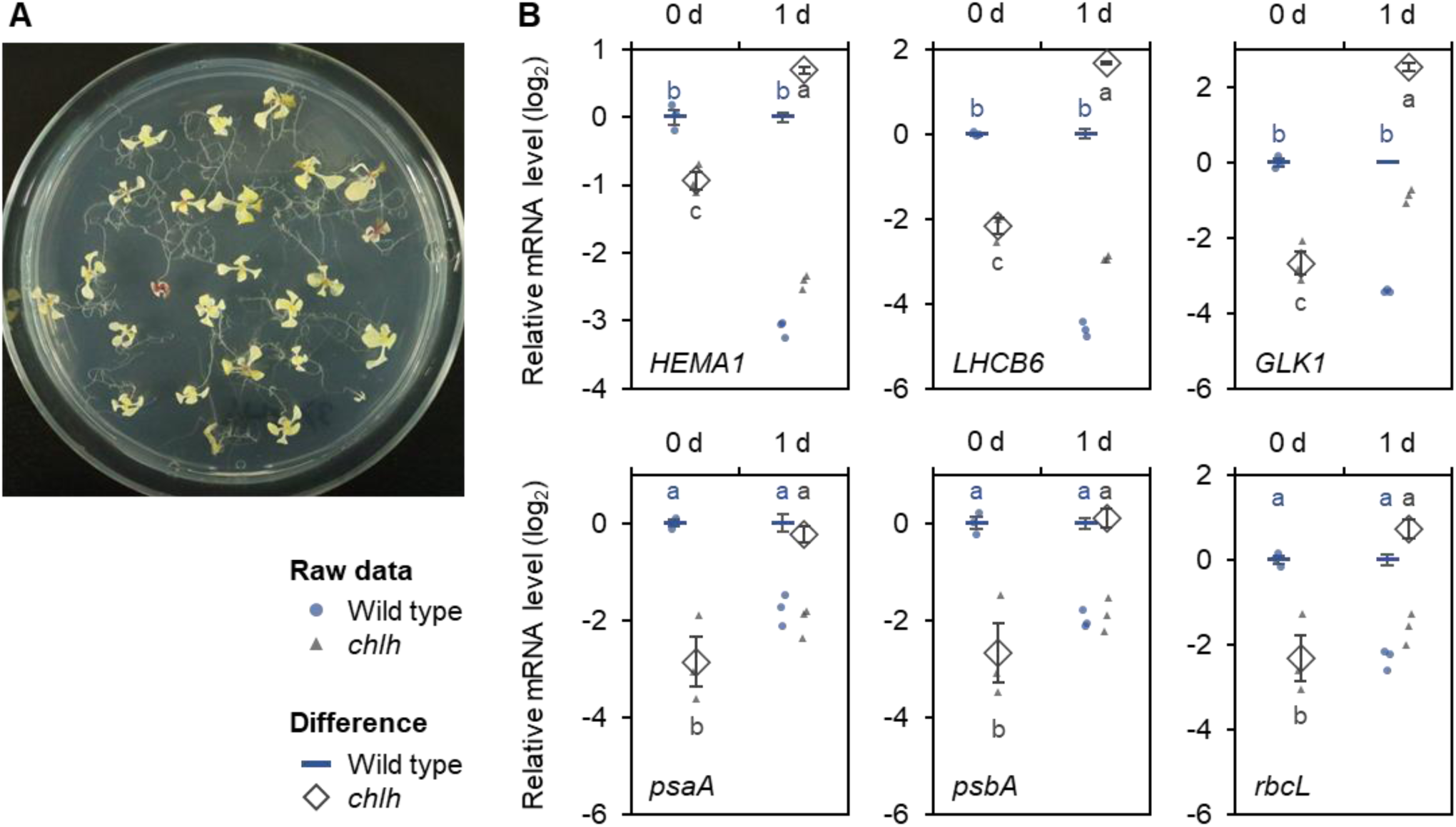
Expression of photosynthesis-related genes in the *chlh* mutant. (A) The visible phenotype of *chlh* seedlings grown for 17 d under continuous light. (B) RT-qPCR analysis of photosynthesis-associated genes. Wild type and *chlh* seedlings grown for 13 and 16 d, respectively, were incubated under continuous light of ∼30 µmol m^− 2^ s^− 1^ or in darkness for 24 h. Transcript levels are presented as the difference from the wild type after normalizing to *ACTIN8* (closed circles and triangles). Then the transcript level of each gene was normalized to the mean value of the corresponding gene in wild type (horizontal bars and open diamonds). Data are means ± SE from three biological replicates. Different letters indicate significant differences (*P* < 0.05, Tukey-Kramer multiple comparison test after two-way ANOVA).

Upon re-illumination after dark incubation for 24 h, the transcript level of nuclear-encoded *HEMA1* slightly increased until 3 h as same as that in *pgp1-2*, whereas the level of *LHCB6* transcript was almost unchanged during the illumination (Fig. 7). The change of mRNA levels of plastid-encoded genes in *chlh* was similar to those in *pgp1-2*, but the light-dependent induction was milder particularly in *rbcL*. Of note, the mRNA levels of *HEMA1, LHCB6*, and *rbcL* in *chlh* were constantly higher than those in *pgp1-2* during illumination.

### Overexpression of *GLK1* upregulates PhANGs but not plastid-encoded genes in *pgp1-2*

GLK transcription factors are known to play a central role in the induction of many PhANGs (Waters *et al.*, 2009). We compared the list of downregulated genes in *pgp1-2* with the list of genes induced by GLK1 and GLK2 (Waters *et al.*, 2009). 50% and 67% of genes induced by GLK1 and GLK2, respectively, were downregulated in *pgp1-2* (Fold change < 0.5, *P* < 0.05; Supplementary Table S4).

Since the mRNA level of *GLK1* was strongly decreased in *pgp1-2* (Figs 5B, 6), we hypothesize that the decreased *GLK1* expression causes downregulation of photosynthesis genes in *pgp1-2*. To test this possibility, we generated the *pgp1-2* mutant overexpressing *GLK1* (*pgp1-2 GLK1*ox) by crossing *pgp1-2* with the 35S promoter-*GLK1* transgenic line (Waters *et al.*, 2008).

The mRNA level of *GLK1* in *pgp1-2 GLK1*ox increased to 300-fold of the wild type (Fig. 9A). The *pgp1-2 GLK1*ox line had more dwarf leaves and shorter petioles than *pgp1-2* (Fig. 9B). The double line contained 3-times higher chlorophyll content with lower chlorophyll *a*/*b* ratio and one-third lower maximum PSII quantum yield (Fv/Fm) than the *pgp1-2* single line (Fig. 9C–E).

**Fig. 9.**
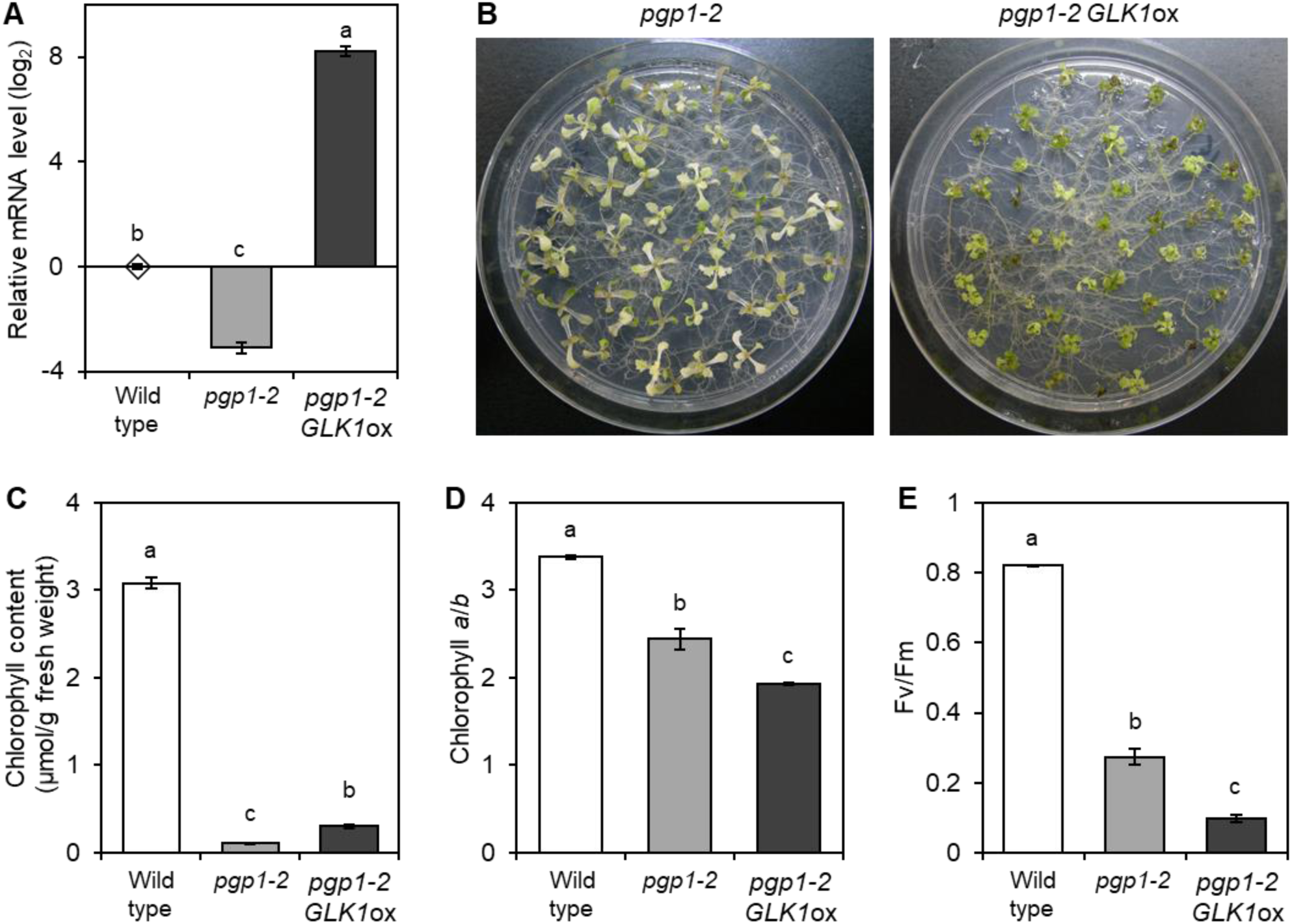
Overexpression of *GLK1* in *pgp1-2*. (A) RT-qPCR analysis of *GLK1* in wild type, *pgp1-2*, and *GLK1* overexpressing *pgp1-2* (*pgp1-2 GLK1*ox). Transcript levels are presented as the difference from the wild type after normalizing to *ACTIN8*. Data are means ± SE from three biological replicates. (B) Visible phenotype of *pgp1-2* and *pgp1-2 GLK1*ox. (C) Chlorophyll accumulation in seedlings. (D) Chlorophyll *a*/*b* ratio. In (C) and (D), data are means ± SE from four (wild type) or five (others) biological replicates. (E) Maximum quantum yield of PSII. Data are means ± SE from five (wild type), seven (*pgp1-2*), and ten (*pgp1-2 GLK1*ox) biological replicates. In all analysis, seedlings grown for 7 d (wild type) and 14 d (*pgp1-2* and *pgp1-2 GLK1*ox) were incubated under 15 µmol photons m^− 2^ s^− 1^ for 7 d. Different letters in (A), (C), (D), and (E) indicate significant differences (*P* < 0.05, Tukey-Kramer multiple comparison test).

We analyzed the mRNA levels of nuclear- and plastid-encoded photosynthesis-related genes in the *pgp1-2 GLK1*ox line (Fig. 10). Among five chlorophyll biosynthesis genes tested, *HEMA1, PORA*, and *PORB* were strongly upregulated by *GLK1* overexpression, whereas *CHLH* and *CHL27* encoding the membranous subunit of Mg-protoporphyrin IX methylester cyclase were not. The mRNA levels of three *LHC* genes, *LHCA4, LHCB4*, and *LHCB6*, were also higher in *pgp1-2 GLK1*ox than in *pgp1-2*. Of two sigma factor genes whose mRNA level was decreased in *pgp1-2, SIG4* was further decreased whereas *SIG1* was not significantly changed by *GLK1* overexpression. Meanwhile, the decreased expression of plastid-encoded genes in *pgp1-2* was not recovered by *GLK1* overexpression. In addition, the mRNA level of *psaA* and *rpoB* even decreased in the double line.

**Fig. 10.**
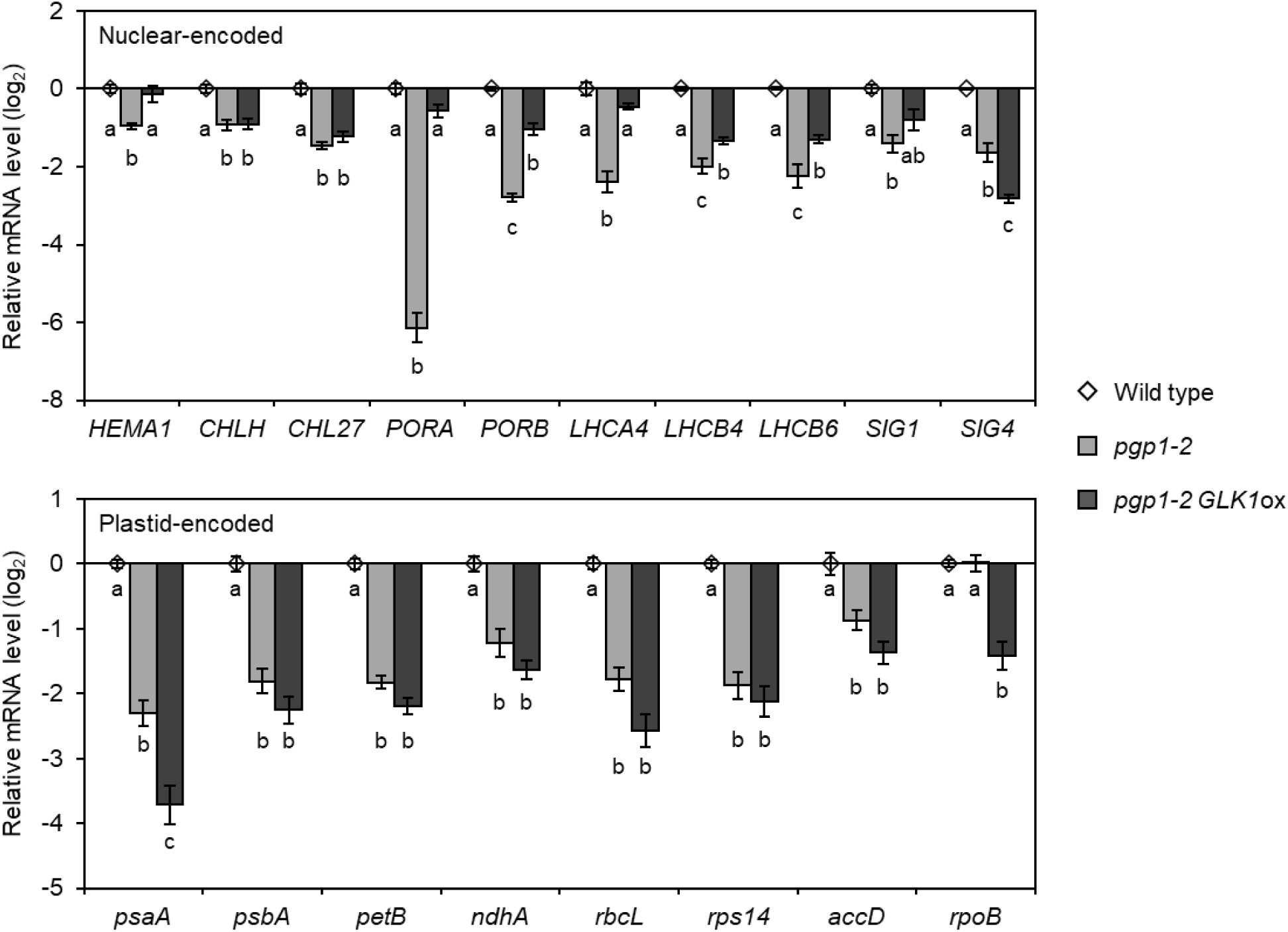
Effects of *GLK1* overexpression on the expression of photosynthesis-associated genes in *pgp1-2*. Seedlings grown for 7 d (wild type) and 14 d (*pgp1-2* and *pgp1-2 GLK1*ox) were incubated under 15 µmol photons m^− 2^ s^− 1^ for 7 d. Transcript levels are presented as the difference from the wild type after normalizing to *ACTIN8*. Data are means ± SE from three biological replicates. Different letters indicate significant differences (*P* < 0.05, Tukey-Kramer multiple comparison test).

## Discussion

### Growth retardation of *pgp1-2* is independent of light-derived stresses

Although PG accounts for only ∼10 mol% of total thylakoid lipids, lack of plastid PG biosynthesis in the *pgp1-2* mutant results in photosynthetic dysfunction, loss of chlorophyll and thylakoid membranes, disrupted leaf morphology, severe growth defects, and enhanced leaf cell death (Fig. 2B, C) (Hagio *et al.*, 2002; Babiychuk *et al.*, 2003; Kobayashi *et al.*, 2015, 2016*a*). To understand how PG deficiency causes such defects during seedling growth, we characterized the *pgp1-2* mutant in detail. Because the *pgp1-2* mutant showed enhanced mRNA expression of ROS-related genes in addition to strong downregulation of PhANGs (Figs. 2A, 3; Supplementary Figs S2, S3), we hypothesized that ROS generated due to light irradiation on disrupted photosynthetic machinery caused the impairment of chloroplast development and morphology in the mutant. However, the impaired leaf development of the *pgp1-2* mutant was not recovered during the growth under very low light (5 µmol photons m^− 2^ s^− 1^) (Fig. 5A). Moreover, TBARS levels used as an oxidative stress marker were not significantly increased in *pgp1-2* even after the treatment of strong red light (Fig. 2E), which was shown to cause strong photoinhibition in this mutant (Kobayashi *et al.*, 2016*a*). Therefore, defects in leaf development may be independent of light exposure or low light may be sufficient to cause impaired leaf development in this mutant.

Of note, the additional mutation of *ex1* in *pgp1-2* did not attenuate the impaired growth (Fig. 2D). EX1 mediates singlet oxygen-induced cell death and the *ex1* mutant fails to induce cell death in response to singlet oxygen generated from chlorophyll or its precursors (Kim and Apel, 2013). Thus, the disrupted leaf morphology in *pgp1-2*, which is accompanied by severe disruption of mesophyll cells inside (Kobayashi *et al.*, 2015), would be independent of singlet oxygen-induced cell death via EX1. This assumption was consistent with our transcriptome data that the *pgp1-2* mutation did not specifically induce singlet oxygen-responsible genes (Fig. 2A). Because additional loss of sulfoquinovosyldiacylglycerol, another major anionic lipid in plastids, in *pgp1-2* causes more severe disruption of leaf development as well as impairments of root and embryo development (Yoshihara *et al.*, 2021), plastid anionic lipids may have a fundamental role in development and morphogenesis of plant organs including leaves.

Meanwhile, the impaired chlorophyll accumulation in *pgp1-2* was partially recovered by *GLK1*ox with increased expression of some genes involved in chlorophyll biosynthesis and light-harvesting (Figs 9, 10). The data suggest that the loss of chlorophyll in this mutant is in part caused by downregulation of these GLK1-targeted genes. By contrast, the seedling growth of *pgp1-2* was not improved and the photochemical efficiency of PSII was even decreased by *GLK1*ox (Fig. 9B, E). PG is a structural and functional component of PSII and essential for its photochemical reaction (Hagio *et al.*, 2000; Sato *et al.*, 2000; Gombos *et al.*, 2002; Sakurai *et al.*, 2003, 2007; Domonkos *et al.*, 2004; Guskov *et al.*, 2009; Umena *et al.*, 2011; Kobayashi *et al.*, 2016*b*,*a*). Therefore, the enhanced chlorophyll accumulation and antenna development with dysfunctional PSII reaction centers by the constitutive PG deficiency would further decrease the photochemical efficiency of PSII in *pgp1-2 GLK1*ox.

### Regulation of PhANGs in *pgp1-2*

Regardless of the growth light intensity, the *pgp1-2* mutant constantly showed lower mRNA expression of photosynthesis-associated genes compared with wild type (Fig. 5B). As a result, the mutant-to-wild type ratio of mRNA levels of these genes was almost unchanged under different light intensities. The data suggest that the *pgp1-2* mutant can upregulate photosynthesis-associated genes in response to increased light intensity as wild type does, while at the same time downregulating these genes at a certain rate. Of note, the transcript levels of some photosynthesis-associated genes were similar between the wild type and *pgp1-2* after the incubation in complete darkness (Fig. 6). Therefore, the suppression of the photosynthesis-associated genes in *pgp1-2* may be dependent on light but independent of its intensity at least in the range examined.

Light response analysis of photosynthesis-associated genes after 1-d dark incubation revealed that the mRNA levels of PhANGs increased during the first 3 h of illumination in *pgp1-2*, although the change was much smaller than in wild type (Fig. 7). The data suggest that light upregulates these genes quickly after illumination even in *pgp1-2*, but it would also trigger a signal to downregulate these genes. Because the *chlh* mutant completely lacking chlorophyll biosynthesis also showed lower mRNA expression of these genes than wild type (Figs 7, 8), impaired chloroplast development under light, whatever it was originally caused by lack of PG biosynthesis or chlorophyll biosynthesis, would be sensed by the plant cell and resulted in downregulation of PhANGs. Of note, a decrease in the *LHCB6* transcript level after prolonged illumination was only observed in *pgp1-2* but not in wild type and *chlh*, the loss of PG may have a stronger impact on downregulation of PhANGs than the chlorophyll deficiency.

Previous studies suggest that GLK1 plays an important role in downregulating PhANGs in response to chloroplast dysfunction; the mRNA level of *GLK1* is decreased by chloroplast dysfunction, which subsequently causes decreased expression of *GLK1* target genes including many PhANGs (Kakizaki *et al.*, 2009; Waters *et al.*, 2009). Consistent with previous studies, mRNA levels of *GLK1* remarkably decreased in *pgp1-2* as well as *chlh* under light, which was accompanied by decreased expression of *GLK1* targets such as genes for chlorophyll biosynthesis and LHC proteins (Figs 6, 8). Because the *GLK1* overexpression in *pgp1-2* partially recovered the decreased expression of PhANGs except for *CHLH, CHL27*, and *SIG*s (Fig. 10), *GLK1* would be one of the key factors determining expression levels of PhANGs in *pgp1-2*. Nonetheless, the expression levels of these GLK1 target genes were not fully recovered in the *pgp1-2 GLK1*ox line despite more than a 200-fold increase of *GLK1* mRNA levels as compared with wild type (Figs 9A, 10). Because the GLK1 protein is degraded in response to chloroplast dysfunction by the ubiquitin-proteasome system (Tokumaru *et al.*, 2017), the *pgp1-2* mutation may induce the degradation of the GLK1 protein and thereby partially cancel the effect of *GLK1* overexpression.

Whereas the *GLK1* expression was suppressed both in *pgp1-2* and *chlh* under light, it was increased after 1-d dark, contrasting with the strong downregulation of *GLK1* in the dark in wild type (Figs 6, 8B). Although the mechanism for the increased *GLK1* expression in the dark-incubated mutants is unknown, the high *GLK1* expression may contribute to relatively higher expression of its target genes such as *LHCB6* in these mutants after 1-d dark treatment.

### Regulation of plastid-encoded genes in *pgp1-2*

Transcriptome analysis of the *pgp1-2* mutant suggests that PEP-dependent transcription is suppressed in this mutant (Fig. 4A). Since the mRNA levels of all *rpo* genes were not altered in *pgp1-2* (Fig. 4A), the impairment of PEP activity in the mutant is not caused by the decreased expression of the PEP core components. The transcript level of *RPOTmp* was upregulated in *pgp1-2* (Fig. 4B), probably as a compensatory effect, which might contribute to maintaining the transcript level of *rpo* genes and other class 3 genes. Although the expression of two *SIG* genes, *SIG1* and *SIG4*, was downregulated in *pgp1-2* (Fig. 4B), these proteins have limited contribution to PEP activity and chloroplast development (Woodson *et al.*, 2013). By contrast, the expression of *SIG2* and *SIG6*, which have a central role in PEP activity in seedlings, was not largely changed in the mutant. Although some nucleoid-associated protein genes were downregulated in *pgp1-2* (Fig. 4C), none of these genes were known to be directly involved in PEP function (Pfalz and Pfannschmidt, 2013). Altogether, our findings suggest an importance of PG in the PEP activity, although the effect of PG deficiency on the protein levels of PEP complexes is an interesting topic for future studies.

We previously reported that the structure of the plastid nucleoid is affected in *pgp1-2* (Kobayashi *et al.*, 2015). Microarray analysis revealed that the mRNA level of three plastid nucleoid-associated genes, i.e., *pTAC8, pTAC16*, and *MFP1*, were downregulated in *pgp1-2* (Fig. 4C). Although the contribution of these proteins on the structural organization of nucleoids remains unknown (Pfalz and Pfannschmidt, 2013; Melonek *et al.*, 2016), this result infers that the downregulation of these factors may be involved in morphological change of plastid nucleoids in *pgp1-2*. It is interesting to note that all three downregulated factors are associated with the plastid membrane (Jeong *et al.*, 2003; Ingelsson and Vener, 2012; Armbruster *et al.*, 2013). Considering that the plastid nucleoid is anchored to the membrane (Krupinska *et al.*, 2013), the downregulation of membrane-associated factors may alter the interaction between the membrane and the nucleoid in *pgp1-2* and subsequently changed the nucleoid structure in plastids.

Wild type and *chlh*, but not *pgp1-2*, showed similar mRNA levels of plastid-encoded genes after 1-d dark incubation (Fig. 8B). In *chlh*, the mRNA levels of plastid-encoded genes under light were similar to those in the dark (Fig. 8B). Together with the data in Fig. 7, these results indicate that the basal expression of plastid-encoded genes in the dark is not inhibited in *chlh*, but induction of these genes by light is suppressed. We previously reported that mRNA levels of *psaA* and *psbA* in light-grown seedlings are not substantially decreased in the *chlh* mutant compared with wild type (Kobayashi *et al.*, 2013*a*), inconsistent with the result in this study. In the previous study, we grew Arabidopsis seedlings in liquid media with rotation in a flask. One possibility is that the growth in liquid media was stressful and affected chloroplast development and photosynthetic gene expression, as suggested by relatively lower chlorophyll content even in wild type (Kobayashi *et al.*, 2013*a*). This specific condition might partly attenuate the light induction of plastid-encoded genes and thus reduce the difference in expression levels of these genes between wild type and the *chlh* mutant.

In contrast to *chlh, pgp1-2* showed lower mRNA levels of *psaA* and *psbA* than wild type even in the dark (Fig. 6). Thus, PG biosynthesis in plastids is required for the basal expression of these genes in the dark in addition to their light induction. Considering that the expression of PEP-transcribed genes was globally downregulated in the *pgp1-2* mutant (Fig. 4A), PG may be constitutively required for the PEP activity in plastids. Because *GLK1* overexpression and chlorophyll accumulation did not recover plastid gene expression in *pgp1-2* (Figs 9, 10), the downregulation of plastid-encoded genes in *pgp1-2* would be independent of impaired chlorophyll biosynthesis and the decreased expression of GLK1 target genes. Moreover, *GLK1* overexpression decreased mRNA levels of *rpoB* in *pgp1-2* (Fig. 10), which might lead to a decreasing trend of plastid-encoded photosynthesis genes. The mechanism of how *GLK1*ox suppresses *rpoB* expression in *pgp1-2* is unknown. We previously reported that the *GLK1* overexpression increases mRNA level of *rpoB* along with *psaA* and *psbA* in wild-type Arabidopsis roots (Kobayashi *et al.*, 2013*b*). An obvious difference between chlorophyll-accumulating roots and *pgp1-2* with *GLK1* overexpression is the functionality of chloroplasts. In *pgp1-2 GLK1*ox, the excess accumulation of chlorophyll over functional photosynthetic proteins and/or the further impairment of the PSII function may lead to the stronger downregulation of plastid-encoded genes.

## Conclusion

Loss of PG in plastids causes the global downregulation of photosynthesis-associated genes encoded in nuclear and plastid genomes and the upregulation of ROS-responsive genes. The downregulation of photosynthesis genes in *pgp1-2* depends on light illumination but independent of the light intensity. Moreover, the developmental defects and the decreased expression of photosynthesis genes in the mutant may occur independently of light-induced oxidative stresses. The decreased PhANG expression in *pgp1-2* is in part caused by the suppression of *GLK1* presumably in a manner related to the loss of chlorophyll accumulation. On the other hand, the downregulation of plastid-encoded genes, particularly those transcribed by PEP, in the mutant is independent of GLK1 and chlorophyll accumulation and suggests a specific involvement of the PG in PEP function.

## Acknowledgements

We thank Shu-Jen Chou (Genomic Technology Core Lab, Institute of Plant and Microbial Biology, Academia Sinica, Taiwan) for technical help with microarray analysis, Nobuyoshi Mochizuki (Graduate School of Science, Kyoto University, Japan) for supplying the *chlh* mutant, Klaus Apel (Boyce Thompson Institute, NY, USA) for providing the *ex1* mutant, and Jane A. Langdale (Department of Plant Sciences, University of Oxford, UK) for the *GLK1*ox line. This work was supported by Grants-in-Aid for Scientific Research (JSPS KAKENHI grant no. 16J10176 and 19J01779 to SF, and grant no. 20K06691 and 18H03941 to KK)).

## Author contributions

SF, KK, and HW: conceptualization; SF, KK, and YN: experimental design; SF and KK: performing experiments; SF, YCL and YL: performing bioinformatic analysis; SF, KK, YCL, YL, YN, and HW: writing.

